# Profiling direct mRNA-microRNA interactions using synthetic biotinylated microRNA-duplexes

**DOI:** 10.1101/005439

**Authors:** Shivangi Wani, Nicole Cloonan

## Abstract

MicroRNAs (miRNAs) are predominantly negative regulators of gene expression that act through the RNA-induced Silencing Complex (RISC) to suppress the translation of protein coding mRNAs. Despite intense study of these regulatory molecules, the specific molecular functions of most miRNAs remain unknown, largely due to the challenge of accurately identifying miRNA targets. Reporter gene assays can determine direct interactions, but are laborious and do not scale to genome-wide screens. Genomic scale methods such as HITS-CLIP do not preserve the direct interactions, and rely on computationally derived predictions of interactions that are plagued by high false positive rates. Here we describe a protocol for the isolation of direct targets of a mature miRNA, using synthetic biotinylated miRNA duplexes. This approach allows sensitive and specific detection of miRNA-mRNA interactions, isolating high quality mRNA suitable for analysis by microarray or RNAseq.

## INTRODUCTION

MicroRNAs (miRNAs) are crucial components of the regulatory circuitry of cells, controlling key biological processes in a broad range of eukaryotic cells. Mature miRNAs are short (∼22nt), and can derive from either the introns of protein-coding genes or from processing of non-coding RNAs through the enzymatic actions of Drosha^1^ and Dicer^2^. The predominant function of miRNAs is to act as guide molecules in the <u>R</u>NA-<u>i</u>nduced <u>s</u>ilencing <u>c</u>omplexes (RISC) and target regions of complementarity in the mRNAs^3^. This targeting inhibits the translation of mRNAs^4–9^ and in some cases leads to mRNA destabilization and degradation^4,10^. Despite intense study of these regulatory molecules, the specific biological and molecular functions of most miRNAs have not yet been identified, largely because our understanding of miRNA-mRNA interactions is far from complete.

Most studied interactions between miRNAs and mRNAs occur in the 3’ UTR^11^, although functional inhibition can occur in the coding region of the mRNA^12–14^, and in the 5’ UTR^15,16^, suggesting that interactions can occur along the entire length of the transcript. Empirical interaction rules have been derived from a small number of experimental validations, and from comprehensive mutation analysis of a small number of miRNAs^17,18^. These studies demonstrate that 6-8nt in the 5’ end of the miRNA (often referred to as the “seed” region) are important, and sometimes sufficient, for binding to mRNAs. Recently, interactions between nucleotides 4 and 15 of the miRNA (centered sites) have also been shown to occur endogenously and mediate mRNA cleavage^19^. The combinatory potential of binding interactions make predicting miRNA targets challenging (especially considering that mismatches, G:U wobble, and bulges can be tolerated), and many programs attempt to restrict the output by considering only interactions driven by a seed, interactions conserved across species, and interactions within the 3’UTR only. Even with these restrictions, computational methods to predict miRNA targets generally perform poorly in experimental validation experiments due to high false positive rates (estimated to be at least 20-40%)^18,20,21^.

Commonly used experimental methods of determining miRNA::mRNA target interactions (such as reporter-gene constructs^22^ and mutation studies^17,18^) are laborious and expensive and cannot scale to whole-genome approaches. Other methods that rely on the degradation of target mRNAs, such as microarray profiling^23^ (in which an exogenous miRNA is introduced to cells, changing mRNA levels that are detected by microarrays), or Parallel Analysis of RNA Ends (PARE)^24^ (in which the cleavage sites of miRNAs are captured by the addition of a 5’ adaptor, prior to standard library preparation for massive-scale sequencing), both miss a substantial fraction of mammalian targets of which the mRNA is not destabilized. Microarray analysis also suffers from the inability to distinguish between the primary effects of miRNA modulation, and downstream effects of changing mRNA target levels. As there is a significant bias towards transcription factors amongst miRNA targets^20^, and because transcription factors act directly and rapidly to alter the level of mRNAs in a cell, distinguishing true miRNA targets from these secondary effects is also a laborious task. Other genome-wide protocols such as HITS-CLIP^25^ or PAR-CLIP^26^, which co-purify the RNA associated with members of RISC, do not preserve the individual relationships between a miRNA and its target, instead relying on computational predictions of miRNA-target interactions, and suffering the same level of false positives described above.

Biotinylated siRNAs were frequently used to isolate and identify the protein components of RISC^27–31^, and some groups have used this approach to capture the direct mRNA targets associated with biotinylated-miRNA duplexes^15,32,33^. Although this method could identify direct targets of miRNAs, the protocol suffered from low specificity^34^ and poor enrichment of targets (an average fold change of ≤ 2)^15^. A modified method, tandem affinity purification of target mRNAs (TAP-Tar), sought to address this by adding a prior isolation step through flag-tagged RISC components^34^. Although the specificity of the pull-down was greatly increased, it still required the use of exogenous, over-expressed proteins in addition to synthetic, modified miRNAs, which is not ideal. Additionally, there are three major concerns that are typically (and correctly) voiced about the original procedure (reviewed by Thomas et al^35^): (i) that the original miR-10a pull-down^15^ did not enrich for known or predicted targets; (ii) that dramatic over-expression could alter the stoichiometry of the detected miRNA-mRNA interactions; and (iii) that dramatic over-expression of a single miRNA could alter the transcriptional network of the cells, leading to RNA changes that could confound our results.

Here we present an improved protocol using biotinylated-miRNA duplexes to detect direct mRNA targets. There are notable differences between the protocol used by Ørom^15,33^, and the one we have used: (i) we use a low concentration of miRNA duplex in a large number of cells to avoid altering the stoichiometry binding site detection; (ii) we have changed the lysis steps from a hypertonic lysis followed by detergent solubilization to a hypotonic lysis step followed by a freeze-thaw; (iii) we have changed the bead type to obtain a higher yield of RNA; (iv) we have blocked the magnetic beads with yeast tRNA to prevent non-specific association of abundant RNA species directly with the beads; and (v) we have introduced several highly stringent wash steps to remove as much non-specific binding as possible. We demonstrate that with these changes, we are able to enrich for known and predicted targets of miRNAs. This protocol does not require exogenous protein expression, but is highly sensitive and specific, and can be used in any cell line (or cell model) where reasonably efficient transient transfection is possible.

## EXPERIMENTAL DESIGN

### Design of miRNA duplexes

There are a few important considerations in the design of the biotin labeled miRNA duplex (**Figure 1**). Biotin should be attached to the 3’ OH group of the mature strand via a C6 linker (available from Integrated DNA Technologies). The exact linker length may not be crucial, as C7 linkers have also been reported to work well^33^. Biotin labeling of the 5’ end will prevent the miRNA from incorporating into RISC, as the 5’ phosphate of the mature strand is required for RISC loading^36,37^. There is some evidence to suggest that overhangs at the 3’ end of each strand (2nt each) are required for unwinding of the double-stranded molecule^38^, and that mismatches in the passenger strand with respect to the mature strand will favor incorporation of mature strand into RISC^39^. Incorporation of additional mismatches through the passenger strand does not appear to affect the performance of the duplex, and may improve the disassociation of the two strands^40^. HPLC purification of the strands prior to annealing is preferred to ensure that full length mature strands are incorporated into RISC.

**Figure 1.**
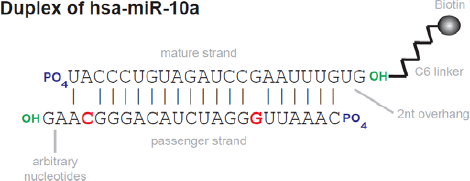
Schematic of miRNA duplex design. An example miRNA duplex is shown for hsa-miR-10a. The mature strand is labeled with a biotin at the 3’ hydroxyl group (green) via a C6 linker. Both the passenger and mature strands should have a 5’ phosphate group (blue). Mismatches between the passenger strand and the mature strand (red) facilitate unwinding of the duplex and incorporation into RISC. The mismatch at position 2 (mature strand) is required; further mismatches are optional.

### Choice of cell line

This protocol should be performed in a cell line that can be easily and reproducibly transfected. Ideally, the transfection efficiency should be at least 30% to minimize the tissue culture required. Many factors affect the successful transfection of mammalian cells (refs), and these parameters will need to be optimized for each cell line. In our experience, conditions that have been optimized for siRNA experiments should work well for miRNA duplexes. It is critical that the cell lines used are Mycoplasma free to ensure reproducible transfection efficiencies. This protocol has been successfully applied in HEK293T, HeLa, and MCF7 cell lines. Whilst the quantitative data gained from this experiment is relevant only in the cell line of origin, qualitative data (such as miRNA-X interacting with mRNA-Y) may be transferable across different cell lines, and can be confirmed by orthogonal methods.

### Concentration of the miRNA duplex

As with any protocol that uses exogenous agents to study endogenous systems, there is a trade-off between efficiency of the protocol and the desire to minimize alterations to the established genetic networks. The ideal concentration of the duplex to be transfected should be optimized for each miRNA, to ensure that known phenotypic effects of miRNA over-expression are absent (although these effects are not always known in advance). We have found that a concentration of 7nM appears to have little effect for the miRNAs we have studied so far (**Figure 2**), and would be a good starting concentration for many miRNAs.

**Figure 2.**
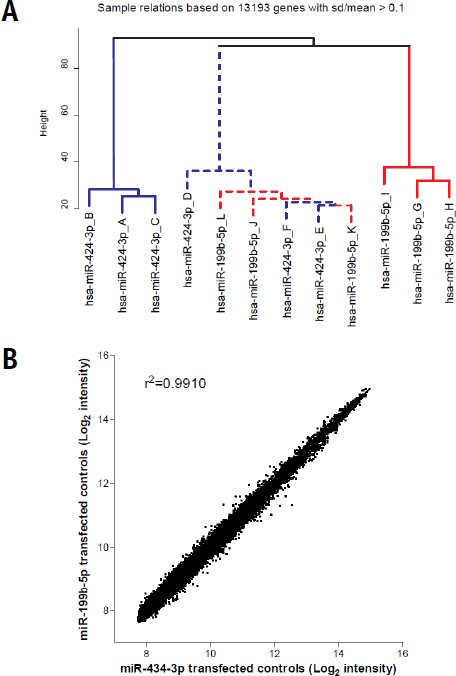
Clustering of microarray samples from two different miRNA pull-downs. (A) Two different miRNA duplexes, hsa-miR-424-3p (blue) and hsa-miR-199b-5p (red), were transfected into HEK293T cells, independently replicated three times. Total RNA samples (dotted lines) and pull-down miRNA enrichments (solid lines) were assayed by microarray, and clustered using the plotSampleRelations function of lumi. There is a very close relationship between the total RNA samples, even though they have been transfected with two different miRNAs. This demonstrates that there is very little effect of either 7nM duplex in this cell line, and that there is no major disruption of the underlying genetic networks upon transfection at this concentration. (B) Correlation of HEK293T control RNA transfected with either miR-424-3p or miR-199b-5p miRNAs. This demonstrates that there is very little effect of either duplex in this cell line, and that there is no major disruption of the underlying genetic networks upon transfection at this concentration.

### Controls

Appropriate controls are critical for the analysis of the experiment. The ideal control for examining direct miRNA-mRNA interactions is the RNA from cells transfected with the biotin-duplex, extracted just prior to pull-down. This controls for changes that the cells may undergo upon transfection with the miRNA, which may not be identified by mock-transfected cells alone. However, this approach is not likely to identify mRNA targets that are directly cleaved by miRNA action, and the inclusion of RNA from mock-transfected cells will help to identify this subset of targets.

## MATERIALS

### REAGENTS

- Cell line of choice (e.g. HEK293T) **▲ CRITICAL** Cells must be Mycoplasma free, and transfection efficiency should be at least 30%
- Gibco^®^ Dulbecco’s Modified Eagle Medium (Invitrogen cat. no. 11995-073)
- Gibco^®^ Fetal Bovine Serum (Invitrogen cat. no. 10099-141)
- Gibco^®^ Phosphate Buffer Saline (Invitrogen cat no. 100.10-023)
- Gibco^®^ Opti-MEM ^®^ Reduced Serum Media (Invitrogen cat. no. 3198-088)
- HiPerfect Transfection Reagent (QIAGEN, cat. no.301702)
- Biotin tagged miRNA duplexes (Integrated DNA Technologies)
- Dynabeads ^®^ MyOne Streptavidin C1 (Invitrogen cat. no. 650-01)
- Buffer Kit (Ambion cat.no. AM9010)
- Sigma-IGEPAL ^®^ CA-630 (Sigma Aldrich cat. no. I8896)
- DL-Dithiothreitol (Sigma Aldrich cat. no. D0632)
- Yeast tRNA (Invitrogen cat.no.15401-011)
- Bovine Serum Albumin (BSA) (for molecular biology, powder; Sigma Aldrich)
- SUPERase•In^™^ (Ambion cat. no.AM2694)
- Complete Mini Protease Inhibitor EDTA free (Roche cat.no. 11836170001)
- RNase/DNase free water
- RNeasy kit (Invitrogen cat. no. 74104) **! CAUTION** buffer RLT contains guanidine isothiocyanate which is harmful and not compatible with disinfectants containing bleach.
- Amicon Ultra-0.5 Centrifugal Filter Devices (Millipore)
- Agilent RNA 6000 Pico Kit (Agilent Technologies cat. no. 5067-1513)
- Illumina ^®^ TotalPrep RNA Amplification kit (Ambion cat. no. AMIL1791)
- Agilent RNA 6000 Nano Kit (Agilent Technologies cat. no.5067-1511)
- Illumina HumanHT-12 v4 Expression BeadChip.
- Dry Ice

### EQUIPMENT

- Cell scrapers
- 10cm tissue culture plates (IWAKI)
- Centrifuge capable of spinning 15mL falcon tubes at 1000g
- 15mL Falcons (BD Bioscience cat. no. 352097)
- Rotating mixer
- DynaMag^™^-2 magnet; magnetic separator (Life technologies cat. no. 123-21D)
- Bench top Microcentrifuges; one set at 4°C and one at room temperature, both should be capable of doing at least 10,000g
- Eppendorf LoBind^®^ tubes; 1.5mL, and 2 mL
- Nanodrop 1000 spectrophotometer (Thermo Scientific)
- Agilent 2100 Bioanalyzer (Agilent Technologies)
- Illumina Bead Array Reader.
- Pipettes and Filter tips (10µL, 20µL, 200μl and 1000μl)

### REAGENT SET UP

- **Biotin tagged miRNA duplexes** Resuspend lyophilized miRNA duplexes to a final concentration of 10uM. Make 20µL aliquots and store at −80°C. **▴ CRITICAL** Repeat freeze thaw cycles should be avoided.
- **Bead Wash Buffer** 5mM Tris-Cl pH 7.5, 0.5mM EDTA, 1M NaCl. Once prepared is stable at room temperature for several months.
- **Solution A** 0.1M NaOH, 0.05M NaCl in RNase/DNase free water. Once prepared is stable at room temperature for several months.
- **Solution B** 0.1M NaCl in RNase/DNase free water. Once prepared is stable at room temperature for several months.
- **Bead Blocking Solution** Prepare RNase/DNase free water containing 1ug/µL BSA and 1ug/µL Yeast tRNA. **▲ CRITICAL** This solution must be made fresh on the day of use.
- **Cell Lysis Buffer** Prepare a solution with final concentrations of the following in RNase/DNase free water −10mM KCl, 1.5mM MgCl2, 10mM Tris-Cl pH 7.5, 5mM DTT, 0.5% Sigma-IGEPAL ^®^ CA-630, 60U/ML SUPERase•In and 1x Complete Mini protease inhibitor. **▲ CRITICAL** This solution must be made fresh on the day of use and must be kept on ice at all times.
- **Wash Buffer** Prepare a solution with final concentrations of the following in RNase/DNase free water −10mM KCl,1.5mM MgCl2,10mM Tris-Cl pH 7.5, 5mM DTT, 0.5% Sigma-IGEPAL ^®^ CA-630, 60U/ML SUPERase•In (Ambion), 1x Complete Mini protease inhibitor (Roche) and 1M NaCl. **▲ CRITICAL** This solution must be made fresh on the day of use and must be kept on ice at all times.

## PROCEDURE

**Step 1-6: Transfecting the Biotin tagged miRNA duplexes into cells. (Day 1) ● TIMING 24.5 hrs, 30 mins hands-on**

1. For each miRNA to be tested, seed 1×10^6^ HEK293T cells in 7mL DMEM + 10% FCS per 10 cm plate in each of 4×10cm TC dishes. Optional: Set up 1×10cm dish with 1×10^6^ HEK293T cells as a mock transfection control if desired.
2. In a sterile 15mL falcon tube, dilute 200pmoles of biotin tagged miRNA in 4mL of OPTI-MEM.
3. Add 160µL of HiPerfect transfection reagent to the diluted miRNA duplex and mix well by vortexing.
4. Incubate the sample at room temperature for 10min to allow the transfection complexes to form.
5. Add 1mL of the transfection complexes to each of the four 10 cm dishes drop-wise. Swirl the plate gently to ensure that the complexes are distributed uniformly across the plate.
6. Incubate the cells with the transfection complexes at 37°C and 5% CO_2_ for 24hr.

**Step 7-20: Bead washing and blocking. Bead preparation can be started on the day of transfection (day 1) on a rotating mixer at 4° overnight, or can be done on the day of cell harvesting (day 2) for 2 hr at room temperature.**

**• TIMING 17.5 hrs if started on day 1, 2.5 hrs if started on day 2, 30 mins hands-on**

1. Resuspend Dynabeads ^®^MyOne Streptavidin C1 in its bottle by vortexing. Transfer 100µL (25µL of bead suspension per 10cm transfected plate) to a 2mL LoBind tube.
2. Place the tube with the bead suspension on the DynaMag^™^-2 magnet for 2 minutes.
3. Using a pipette, aspirate and discard supernatant before removing tube from the DynaMag^™^-2 magnets.
4. Add 100µl of bead wash buffer to the beads. Pipette several times to ensure the beads are washed sufficiently.
5. Repeat steps 8 - 10 twice more for a total of three washes.
6. RNase Freeing Beads: After the third wash resuspend the beads in 100µl Solution A. Mix well by pipetting several times. Let beads incubate at room temperature for 2 minutes.
7. Place the tube containing the bead solution on the DynaMag^™^-2 magnet for 2 minutes.
8. Using a pipette, aspirate and discard supernatant before removing tube from the DynaMag^™^-2 magnets.
9. Repeat steps 12-14 once more.
10. Resuspend beads in 100µl Solution B. Mix well by pipetting several times.
11. Place the tube containing the bead solution on the DynaMag^™^-2 magnet for 2 minutes.
12. Using a pipet, aspirate and discard supernatant before removing tube from the DynaMag^™^-2 magnets.
13. Resuspend beads in 200µl Bead blocking solution. Pipette several times to mix.
14. Place the tube containing the bead on a rotating mixer at 4°C overnight if starting on day 1, or allow mixing at room temperature for 2 hrs if starting on day 2.

**Steps 21-36 Harvesting and lysing transfected cells *(Day 2)* ● TIMING 1hr, 30 mins hands-on**

1. Prepare cell Lysis Buffer and keep buffer on ice till required. **▲ CRITICAL** this buffer must be made fresh on the day of use.
2. Retrieve transfection plates from step 6 of the protocol. Aspirate and discard media.
3. To each 10 cm plate, add 2 mL of DMEM + 10% FCS and using a cell scraper gently lift off the cells from the plate.
4. Using a pipette transfer the cells to a 15mL falcon tube. <u>Note:</u> Cells from each of the 4x 10cm plates should be pooled at this stage so your falcon tube should contain 8mL cells.
5. Centrifuge the falcon tubes containing the harvested cells at 1000rpm for 5 minutes at room temperature.
6. Aspirate and discard the supernatant. Add 5mL of PBS to the cell pellet and then pipette gently to wash and resuspend the cells.
7. Centrifuge the falcon tube at 1000rpm for 5 minutes at room temperature.
8. Aspirate and discard supernatant.
9. Add 250µl cell lysis buffer to each pellet and gently pipette to resuspend cells.
10. Put cells in lysis buffer on dry ice for 5 mins.
11. Allow the cells to thaw out at room temperature and then transfer lysate to a 1.5mL LoBind tube. **▲ CRITICAL** this freeze thaw step allows for better lysis of cells.
12. Centrifuge tube at 13,000rpm in a bench top centrifuge set at 4°C for 2 minutes.
13. Transfer the cleared cell lysate to a clean 2mL LoBind tube, leaving behind the soft pellet. The final volume of cleared lysate should be ∼240µl-250µL.
14. Transfer 10µL of this cleared lysate to a clean 1.5mL LoBind tube and keep on ice. This lysate will serve as the control lysate RNA. <u>Note:</u> If processing a mock transfected control, carry out steps 22-33 of protocol on the mock transfection plate and keep mock control lysate on ice for later use.
15. Add NaCl to the cleared lysate to give a final concentration of 1M. For example, add 60µL of 5M NaCl to 240µL of lysate, giving a final concentration of 1M NaCl in 300µL.
16. Place the tube containing cleared lysate and NaCl on ice.

**Steps 37-44 Bead Preparation ● TIMING 15 mins, 15 mins hands-on**

1. Prepare wash Buffer and keep buffer on ice till required. **▲ CRITICAL** This buffer must be made fresh on the day of use.
2. Place the tube with the beads from step 20 on the DynaMag^™^-2 magnets for 2 minutes.
3. Aspirate and discard supernatant using a pipette.
4. Add 100µl of wash buffer and Resuspend beads by pipetting several times.
5. Place tube on the DynaMag^™^-2 magnets for 2 minutes.
6. Aspirate the supernatant using a pipette and discard. Remove the tube from the DynaMag^™^-2 magnets.
7. Repeat steps 40-42
8. Resuspend beads in 300µl of wash buffer.

**Steps 45-52 Target mRNA capture and post capture bead washing ● TIMING 45 mins, 15 mins hands-on**

1. Add the 300µl cell lysate + NaCl from step 36 to the 300µl prepared beads from step 45. Place tubes on a rotating mixer at room temperature and incubate for 30 minutes.
2. Place the tube containing the cell lysate + Beads on the DynaMag^™^-2 magnets for 2 minutes.
3. Aspirate supernatant using pipette and discard. Remove the tube from the DynaMag^™^-2 magnets.
4. Wash beads in 300µl wash buffer. Mix well by pipetting several times.
5. Place the tube containing Beads + wash buffer on the DynaMag^™^-2 magnets for 2 minutes.
6. Aspirate supernatant using pipette and discard. Remove the tube from the DynaMag^™^-2 magnets.
7. Repeat steps 48 - 50 three times. **▲ CRITICAL** These washes are essential and are important for the removal of non-specifically bound products from the beads.
8. Once the washes are complete, resuspend the beads in 100µL of RNase/DNase free water, store on ice and proceed immediately to the next step. **▲ CRITICAL** The target mRNAs are captured on these beads. Do not discard.

**Steps 53-59 Target mRNA and control lysate RNA purification ● TIMING 1hr, 30 mins hands-on**

1. Target mRNAs are purified off the Dynabeads^®^ MyOne Streptavidin C1 using a Qiagen RNeasy kit according to manufacturer’s RNA clean-up protocol. RNA should be eluted twice in 50µl RNase/DNase free water. The final volume of RNA will be 100µl.
2. Alongside the captured target mRNAs also purify the control lysate RNA from step 34 of this protocol. Make up the volume of the control lysate to 100µL by adding 90µL of RNase /DNase free water. Purify the control samples using a Qiagen RNeasy kit according to manufacturer’s RNA clean-up protocol. The control RNA can be eluted in 30-50µL of RNase/DNase free water.
3. Add 400µl RNase/DNase free water to the purified target mRNA to bring the total RNA volume to 500µL. Similarly, add 450µL of RNase/DNase free water to the control RNA to bring it to a total volume of 500µL.
4. Assemble an Amicon Ultra-0.5 Centrifugal filter device into a fresh collection tube. Now transfer the 500µL of target mRNA and the control RNA each into separate Amicon Ultra-0.5 Centrifugal filter devices.
5. Put the Amicon Ultra-0.5 Centrifugal filter devices containing the RNAs into a tabletop centrifuge and spin at 13,000 rpm for 30 minutes at room temperature.
6. Next place the Amicon Ultra-0.5 Centrifugal filter device upside down in a fresh collection tube and re-centrifuge in the tabletop centrifuge at 5000 rpm for 2 minutes at room temperature.
7. The RNAs should have eluted out of the filtration device into the collection tubes in a final volume of approximately 20µl-25µL. <u>Note:</u> If you have a mock control lysate, put it through steps 54 to 58 of the protocol. **▲ CRITICAL** This extra cleanup step is essential to remove carry-over organic contaminants which can inhibit the labeling and amplification of the RNA sample.

**▪ PAUSE POINT** Samples can now be stored at −80°C. Alternatively, continue with quantification of RNA.

**Step 60 Target mRNA and control RNA Quantification ● TIMING 45mins, 20 mins hands-on**

1. The target mRNA and the control lysate RNA can now be quantified on the Nanodrop 1000 spectrophotometer and on an Agilent 2100 Bioanalyser using the Agilent RNA 6000 Pico kit using manufacturer’s protocol.

**▪ PAUSE POINT** Samples can now be stored at −80°C or continue on with RNA amplification.

**Step 61-62 Target mRNA and control RNA Labeling and Amplification ● TIMING 23 hrs, 2hrs hands-on**

1. Amplify and label 50ng of captured target mRNA using Illumina^®^ TotalPrep RNA Amplification kit according to manufacturer’s instructions. Also amplify and label 50ng of control RNA. **▲ CRITICAL** Carry out the 14 hour incubation for the IVT step.
2. Quantify the amplified RNAs on the Nanodrop 1000 spectrophotometer and then check the size distribution on an Agilent 2100 Bioanalyser using the Agilent RNA 6000 Nano kit using manufacturer’s protocol.

**Step 63-65 Hybridization of samples onto Illumina^®^ Human HT-12 array and scanning ● TIMING 18hrs, 2hrs hands-on**

1. Hybridize 750ng of amplified cRNA onto an Illumina^®^ Human HT-12 array following manufacturer’s protocol.
2. Scan the arrays using an Illumina BeadArray Reader.
3. Extract the expression measurements using the GenomeStudio software.

**● TIMING**

Timings for each step are based on processing 1-12 samples for final hybridization onto a single Illumina^®^ Human HT-12 array.

Step 1-6: Transfecting the Biotin tagged miRNA duplexes into cells: 24.5 hrs

Step 7-20: Bead washing and blocking: 17.5 hrs if started on Day1 and 2.5 hrs if started on Day 2

Steps 21-36: Harvesting and lysing transfected cells (Day 2): 1hr

Steps 37-44: Bead Preparation: 15 mins

Steps 45-52: Target mRNA capture and post-capture bead washing: 45 mins

Steps 53-59: Target mRNA and control lysate RNA purification: 1hr

Step 60: Target mRNA and control RNA Quantification: 45 mins

Step 61-62: Target mRNA and control RNA Labeling and Amplification: 23 hrs

Step 63-65: Hybridization of samples onto Illumina^®^ Human HT-12 array and scanning: 18hrs

**? TROUBLESHOOTING**

Troubleshooting advice for possible problems when affinity purifying targets of miRNAs can be found in **Table 1**.

**TABLE 1.**
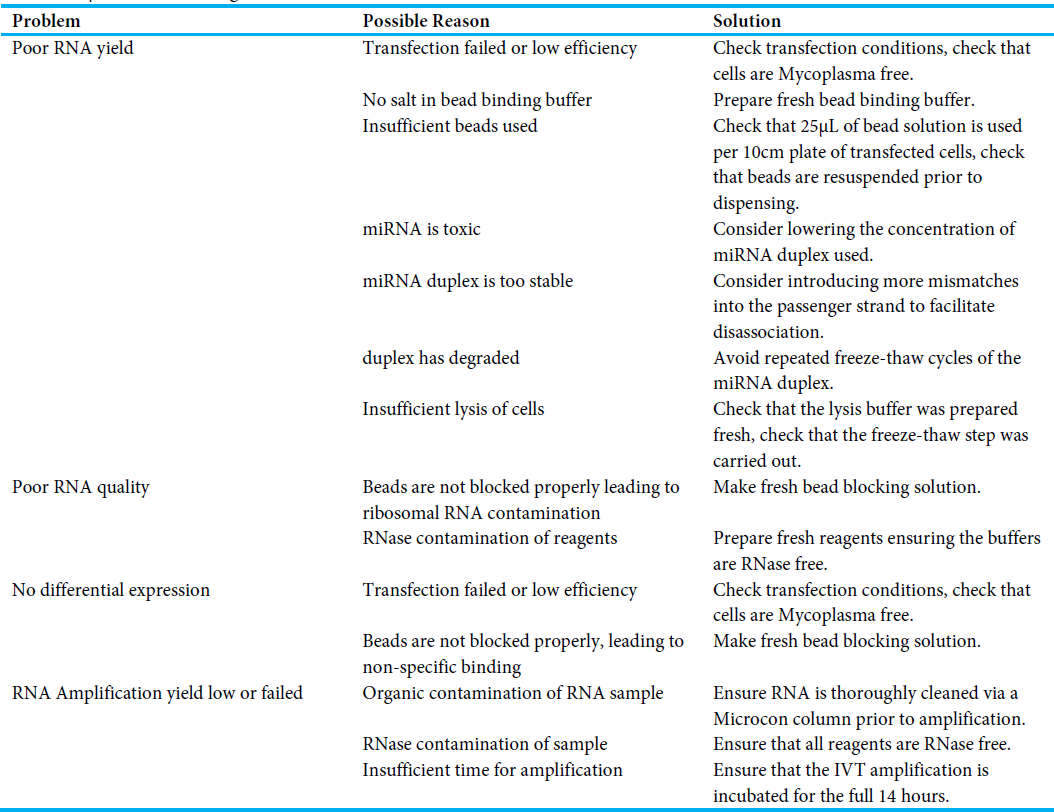
Troubleshooting table.

## ANTICIPATED RESULTS

The amount of RNA that is pulled down in this protocol will vary from experiment to experiment, depending on the miRNA and the cell line used. The minimum amount of RNA suggested by the manufacturer for RNA amplification is 50ng, but we have been able to robustly identify miRNA targets using only 20ng of input. Typical profiles of pulled-down mRNA and amplified mRNA are shown in **Figure 3A** and **B**, respectively.

**Figure 3.**
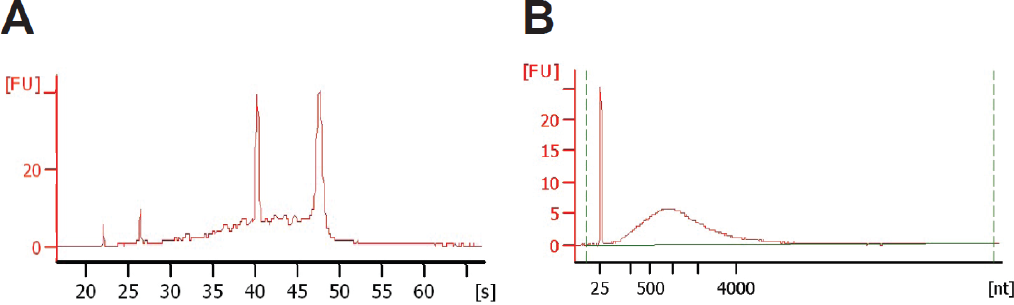
Bioanalyzer profiles of RNA. (A) Typical Bioanalyzer of RNA from hsa-miR-17-5p biotin pull-down after step 59 of the protocol. (B) Typical Bioanalyzer profile of amplified RNA after step 62.

The data from the Illumina microarrays can be analyzed in R using the lumi^41^ and limma^42^ packages. An example R script has been provided to analyze Illumina microarray data starting with the output from GenomeStudio (**Supplementary File 1**). Briefly, arrays should be background corrected, data transformed using a variance-stabilizing transformation^43^, and then normalized (within and between arrays) using the robust spline method^44^. All probes with a detection p-value less than 0.01 in all control samples can be retained for further analysis. The lmFit and eBayes functions in the limma package are used to test for differential expression between the pull-down samples and the control samples for each miRNA. The Benjamini-Hochberg correction is applied in order to account for multiple testing, and probes with an adjusted one-sided p-value less than 0.05 (lower-tailed p-value in the code in Supplementary File 1) are considered to be significantly enriched in the pull-down, and thus targets of the miRNA.

The number of microarray probes (and corresponding genes or transcripts) detected as differentially expressed will vary for each individual miRNA^45^, but hundreds to thousands of enriched targets are not uncommon. Additional stringency can be gained by introducing a fold-change threshold, but we found that applying a threshold of 2-fold change in enrichment did not significantly change either the gene set enrichment analysis, or the cross-validation rate using luciferase assays^40^.

Using the same miRNA duplex that was used by Ørom et al. (hsa-miR-10a)^15^, we performed the pull-down and produced a volcano plot (**Figure 4**, left), which plots the fold change versus the −log_10_ transformed significance of that fold change for every probe on the microarray. We produced a similar plot using the data analyzed by Ørom et al^15^ (ArrayExpress accession number E-MEXP-1375; **Figure 4**, right). Whereas the data from the latter has few genes passing a significance threshold, and a very limited range of differential expression, the data from our protocol shows substantial improvement in all these areas. Additionally, we see the expected enrichment of both known and predicted mRNA targets, confirming the usefulness of this protocol. We have used this approach to determine the direct mRNA targets of more than 10 miRNAs^40,45–47^, to determine that isomiRs are target functionally related pathways as their canonical partners^40^, and to determine that a large proportion of interactions are not mediated by the seed sequence^45^.

**Figure 4.**
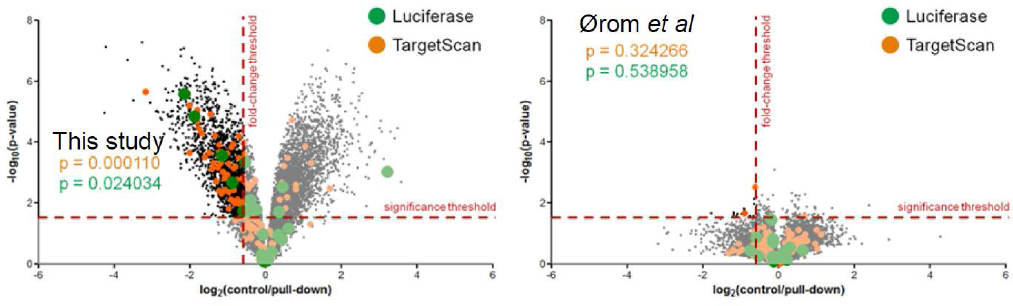
Volcano plots showing the significance of the difference in expression between the indicated pull-down and the mock-transfected control, for all transcripts expressed in control cells. Targets predicted by TargetScan or validated previously via luciferase assay are indicated by orange and green dots, respectively. A comparison between miR-10a biotin pull-down in this study (left) and by Ørom et al^15^ (right). P values for the enrichment of Luciferase validated targets or TargetScan predicted targets are indicated. Red dashed lines indicated the significance threshold and the fold-change threshold used in this study. For the pull-downs performed by Ørom et al., no enrichment of known or predicted targets was observed. For the pull-downs using our improved protocol, we observe substantial increases in dynamic range, and statistically significant enrichment of both known and predicted targets.

## ACKNOWLEDGEMENTS

We are grateful to all the members of QCMG for their assistance, and particularly to Sean M. Grimmond for his patience as we optimized the protocol in his laboratory. We also thank Anita L. Steptoe for tissue culture, Katia Nones for microarray support, Keerthana Krishnan for luciferase assays, and Hilary C. Martin for bioinformatics analysis.

## REFERENCES

1. Han, J. et al. The Drosha-DGCR8 complex in primary microRNA processing. Genes Dev. 18, 3016–27 (2004).

2. Hutvágner, G. et al. A cellular function for the RNA-interference enzyme Dicer in the maturation of the let-7 small temporal RNA. Science (80-.). 293, 834–8 (2001).

3. Bartel, D. P. MicroRNAs:: Genomics, Biogenesis, Mechanism, and Function. Cell 116, 281–297 (2004).

4. Humphreys, D. T., Westman, B. J., Martin, D. I. K. & Preiss, T. MicroRNAs control translation initiation by inhibiting eukaryotic initiation factor 4E/cap and poly(A) tail function. PNAS 102, 16961–6 (2005).

5. Maroney, P. a, Yu, Y., Fisher, J. & Nilsen, T. W. Evidence that microRNAs are associated with translating messenger RNAs in human cells. Nat. Struct. Mol. Biol. 13, 1102–7 (2006).

6. Mathonnet G. et al. MicroRNA inhibition of translation initiation in vitro by targeting the cap-binding complex eIF4F. Science (80-.). 317, 1764–7 (2007).

7. Nottrott, S., Simard, M. J. & Richter, J. D. Human let-7a miRNA blocks protein production on actively translating polyribosomes. Nat. Struct. Mol. Biol. 13, 1108–14 (2006).

8. Petersen, C. P., Bordeleau, M.-E., Pelletier, J. & Sharp, P. a. Short RNAs repress translation after initiation in mammalian cells. Mol. Cell 21, 533–42 (2006).

9. Pillai, R. S. et al. Inhibition of translational initiation by Let-7 MicroRNA in human cells. Science (80-.). 309, 1573–6 (2005).

10. Wu, L., Fan, J. & Belasco, J. G. MicroRNAs direct rapid deadenylation of mRNA. PNAS 103, 4034–9 (2006).

11. Rigoutsos, I. New tricks for animal microRNAS: targeting of amino acid coding regions at conserved and nonconserved sites. Cancer Res. 69, 3245–8 (2009).

12. Shen, W.-F., Hu, Y.-L., Uttarwar, L., Passegue, E. & Largman, C. MicroRNA-126 regulates HOXA9 by binding to the homeobox. Mol. Cell. Biol. 28, 4609–19 (2008).

13. Duursma, A. M., Kedde, M., Schrier, M., le Sage, C. & Agami, R. miR-148 targets human DNMT3b protein coding region. RNA 14, 872–7 (2008).

14. Forman, J. J., Legesse-Miller, A. & Coller, H. a. A search for conserved sequences in coding regions reveals that the let-7 microRNA targets Dicer within its coding sequence. Proc. Natl. Acad. Sci. U. S. A. 105, 14879–84 (2008).

15. Ørom, U. A., Nielsen, F. C. & Lund, A. H. MicroRNA-10a binds the 5’UTR of ribosomal protein mRNAs and enhances their translation. Mol. Cell 30, 460–71 (2008).

16. Lytle, J. R., Yario, T. a & Steitz, J. a. Target mRNAs are repressed as efficiently by microRNA-binding sites in the 5’ UTR as in the 3’ UTR. Proc. Natl. Acad. Sci. U. S. A. 104, 9667–72 (2007).

17. Brennecke, J., Stark, A., Russell, R. B. & Cohen, S. M. Principles of microRNA-target recognition. PLoS Biol. 3, e85 (2005).

18. Lewis, B. P., Burge, C. B. & Bartel, D. P. Conserved seed pairing, often flanked by adenosines, indicates that thousands of human genes are microRNA targets. Cell 120, 15–20 (2005).

19. Shin, C. et al. Expanding the microRNA targeting code: functional sites with centered pairing. Mol. Cell 38, 789–802 (2010).

20. Lewis, B. P., Shih, I., Jones-Rhoades, M. W., Bartel, D. P. & Burge, C. B. Prediction of mammalian microRNA targets. Cell 115, 787–98 (2003).

21. Krek, A. et al. Combinatorial microRNA target predictions. Nat. Genet. 37, 495–500 (2005).

22. Zeng, Y., Wagner, E. J. & Cullen, B. R. Both Natural and Designed Micro RNAs Can Inhibit the Expression of Cognate mRNAs When Expressed in Human Cells. Mol. Cell 9, 1327–1333 (2002).

23. Lim, L. P. et al. Microarray analysis shows that some microRNAs downregulate large numbers of target mRNAs. Nature 433, 769–73 (2005).

24. German, M. a, Luo, S., Schroth, G., Meyers, B. C. & Green, P. J. Construction of Parallel Analysis of RNA Ends (PARE) libraries for the study of cleaved miRNA targets and the RNA degradome. Nat. Protoc. 4, 356–62 (2009).

25. Licatalosi, D. D. et al. HITS-CLIP yields genome-wide insights into brain alternative RNA processing. Nature 456, 464–9 (2008).

26. Hafner, M. et al. Transcriptome-wide identification of RNA-binding protein and microRNA target sites by PAR-CLIP. Cell 141, 129–41 (2010).

27. Ameres, S. L., Martinez, J. & Schroeder, R. Molecular basis for target RNA recognition and cleavage by human RISC. Cell 130, 101–12 (2007).

28. Rand, T. a, Petersen, S., Du, F. & Wang, X. Argonaute2 cleaves the anti-guide strand of siRNA during RISC activation. Cell 123, 621–9 (2005).

29. Chu, C. & Rana, T. M. Translation repression in human cells by microRNA-induced gene silencing requires RCK/p54. PLoS Biol. 4, e210 (2006).

30. Martinez, J., Patkaniowska, A., Urlaub, H., Lührmann, R. & Tuschl, T. Single-stranded antisense siRNAs guide target RNA cleavage in RNAi. Cell 110, 563–74 (2002).

31. Kedde, M. et al. RNA-binding protein Dnd1 inhibits microRNA access to target mRNA. Cell 131, 1273–86 (2007).

32. Lal, A. et al. Capture of MicroRNA–Bound mRNAs Identifies the Tumor Suppressor miR-34a as a Regulator of Growth Factor Signaling. PLoS Genet. 7, e1002363 (2011).

33. Orom, U. A. & Lund, A. H. Isolation of microRNA targets using biotinylated synthetic microRNAs. Methods 43, 162–5 (2007).

34. Nonne, N., Ameyar-Zazoua, M., Souidi, M. & Harel-Bellan, A. Tandem affinity purification of miRNA target mRNAs (TAP-Tar). Nucleic Acids Res. 38, e20 (2010).

35. Thomas, M., Lieberman, J. & Lal, A. Desperately seeking microRNA targets. Nat. Struct. Mol. Biol. 17, 1169–74 (2010).

36. Parker, J. S., Roe, S. M. & Barford, D. Structural insights into mRNA recognition from a PIWI domain-siRNA guide complex. Nature 434, 663–6 (2005).

37. Ma, J.-B. et al. Structural basis for 5’-end-specific recognition of guide RNA by the A. fulgidus Piwi protein. Nature 434, 666–70 (2005).

38. Ma, J.-B., Ye, K. & Patel, D. J. Structural basis for overhang-specific small interfering RNA recognition by the PAZ domain. Nature 429, 318–22 (2004).

39. Schwarz, D. S. et al. Asymmetry in the assembly of the RNAi enzyme complex. Cell 115, 199–208 (2003).

40. Cloonan, N. et al. MicroRNAs and their isomiRs function cooperatively to target common biological pathways. Genome Biol. 12, R126 (2011).

41. Du, P., Kibbe, W. a & Lin, S. M. lumi: a pipeline for processing Illumina microarray. Bioinformatics 24, 1547–8 (2008).

42. Smyth, G. K. Linear models and empirical bayes methods for assessing differential expression in microarray experiments. Stat. Appl. Genet. Mol. Biol. 3, Article3 (2004).

43. Lin, S. M., Du, P., Huber, W. & Kibbe, W. a. Model-based variance-stabilizing transformation for Illumina microarray data. Nucleic Acids Res. 36, e11 (2008).

44. Workman, C. et al. A new non-linear normalization method for reducing variability in DNA microarray experiments. Genome Biol. 3, research0048 (2002).

45. Martin, H. C. et al. Imperfect centered miRNA binding sites are common and can mediate repression of target mRNAs. Genome Biol. 15, R51 (2014).

46. Krishnan, K. et al. MicroRNA-182-5p targets a network of genes involved in DNA repair. RNA 19, 230–242 (2013).

47. Krishnan, K. et al. miR-139-5p is a regulator of metastatic pathways in breast cancer. RNA 19, 1767–80 (2013).

